# LD scores are associated with differences in allele frequencies between populations but LD score regression can still distinguish confounding from polygenicity

**DOI:** 10.1101/562629

**Authors:** Mason Alexander, David Curtis

**Affiliations:** UCL Genetics Institute, UCL, Darwin Building, Gower Street, London WC1E 6BT.; Centre for Psychiatry, Barts and the London School of Medicine and Dentistry, Charterhouse Square, London EC1M 6BQ.

**Keywords:** LD score regression, case-control, association, HapMap.

## Abstract

The LD score regression method tests whether there is an association between the LD score and allele frequency differences between cases and controls. It makes the assumption that there is no association between LD score and allele frequency differences between populations and hence that any observed association is due to a polygenic effect rather than population stratification. This assumption has not previously been tested. In comparisons between HapMap populations we observe that there is indeed an association between the LD score and allele frequency differences. However this effect is small and when we carry out simulations of large case-control samples the effect becomes negligible. We conclude that if the intercept is small then any increase in mean chi-squared does indeed reflect a polygenic effect rather than population stratification.

## Introduction

LD score regression was proposed as a method to distinguish polygenic effects in genome-wide association studies (GWASs) from confounding biases such as cryptic relatedness and population stratification (Bulik-Sullivan *et al*., 2015). In a GWAS one seeks to detect differences in allele frequencies between cases and controls due to variants which either directly affect phenotype or else which are in linkage disequilibrium (LD) with causal variants. A problem is that variant allele frequencies can vary between populations with different ancestries and hence a false positive GWAS signal can be produced if case and control samples are not properly matched. In order to detect that there are variants truly associated with disease risk, LD score regression makes two assumptions. The first is that variants are more likely to be in LD with a causal variant if they are in LD with other nearby variants, as measured by their LD score. Under this assumption, there will be a positive correlation across variants between the LD score and a measure of difference in allele frequency between cases and controls such as the chi-squared statistic. The second assumption is that there will be no association between the LD score and the difference in allele frequency between populations. Under these two assumptions one can perform linear regression of the chi-squared onto the LD score and a positive gradient will indicate a polygenic effect on risk while the intercept will capture the effect of population stratification.

In the original publication the second assumption was tested using Psychiatric Genetics Consortium controls from seven European cohorts and by computing association statistics between pairs of cohorts (Bulik-Sullivan *et al*., 2015). For all pairs of cohorts there was minimal correlation, with the largest R-squared for any pair reported to be 0.000255. However all the cohorts used had been selected to be of white European origin and it is not known whether the correlation might be stronger if more ancestrally diverse cohorts were utilised. Although case-control studies of moderately rare traits will typically use subjects which are intended to be well-matched for ancestry, this may not be the case for studies which use very large samples derived from more diverse sources. For example, a recent study of risk tolerance used a sample of over 900,000 subjects recruited from UK Biobank and 23andMe and reported an LD score intercept of 1.04 and a mean chi-squared of 1.85 (Karlsson Linnér *et al*., 2019). One could speculate that if the chi-squared between different ancestries was correlated with the LD score then if there were a slight enrichment for one ancestry among cases then this could lead to an inflated mean chi-squared which reflected population stratification rather than a true polygenic effect.

## Method

In order to examine the correlation between LD scores and allele frequency differences between populations of different ancestries we used the same HapMap datasets as we had used to demonstrate that the polygenic risk score for schizophrenia was associated with ancestry (Curtis, 2018). The merged post-QC phase I+II and III HapMap (International HapMap 3 Consortium *et al*., 2010) genotype files were downloaded from ftp://ftp.ncbi.nlm.nih.gov/hapmap/genotypes/2010-08_phaseII+III/forward/. The file called *scz2.prs.txt.gz*, containing ORs and p values for 102,636 LD-independent single nucleotide polymorphism markers (SNPs), was downloaded from the Psychiatric Genetics Consortium (PGC) website (www.med.unc.edu/pgc/results-and-downloads). This is the training set produced as part of the PGC2 schizophrenia GWAS (Schizophrenia Working Group of the Psychiatric Genomics Consortium, 2014). This SNP set was obtained from the imputed GWAS genotypes by first excluding uncommon SNPs (MAF < 10%), low-quality variants (imputation INFO < 0.9), indels, and SNPs in the extended MHC region (chr6:25-34 Mb). The SNPs were then LD pruned and “clumped”, by discarding SNPs within 500 kb of, and in r^2^ ≥ 0.1 with, another SNP which was more significantly associated with schizophrenia. Autosomal SNPs were selected if they appeared in this training dataset and if they had also been genotyped in all 11 of the HapMap cohorts, yielding a reduced set of 32,588 LD-independent SNPs. HapMap subjects with genotyping call rate < 0.9 were removed, leaving a sample of 1,397.

After QC, the 11 HapMap cohorts consisted of the following samples: ASW - African ancestry in Southwest USA, N=87; CEU - Utah residents with Northern and Western European ancestry, N=174; CHB - Han Chinese in Beijing, China, N=139; CHD - Chinese in Metropolitan Denver, Colorado, N=109; GIH - Gujarati Indians in Houston, Texas, N=101; JPT - Japanese in Tokyo, Japan, N=116; LWK - Luhya in Webuye, Kenya, N=110; MEX - Mexican ancestry in Los Angeles, California, N=86; MKK - Maasai in Kinyawa, Kenya, N=184; TSI - Toscani in Italia, N=102; YRI - Yoruba in Ibadan, Nigeria, N=209. The set of SNPs was reduced to the 30,753 for which european LD scores were available, as contained in the file https://data.broadinstitute.org/alkesgroup/LDSCORE/eur_w_ld_chr.tar.bz2 obtained from https://github.com/bulik/ldsc/wiki/LD-Score-Estimation-Tutorial. The allele frequencies in the CEU cohort were compared with those in each of the other ten cohorts using the *assoc* function of *plink 1.09beta* to produce a chi-squared statistic for each SNP (www.cog-genomics.org/plink/1.9/) (Purcell *et al*., 2007, 2009; Chang *et al*., 2015). Linear regression of the chi-squared statistics onto the LD scores was carried out using R version 3.3.2 (R Core Team, 2014).

To obtain results for a set of SNPs which were not LD-pruned, the same LD score regression analysis between CEU and the other cohorts was then repeated using all 15,216 chromosome 22 SNPs which were present in HapMap and for which european LD scores were available.

In order to assess the effects of population stratification, datasets were constructed which were intended to reflect varying proportions of CEU and YRI ancestry. A set of 200 controls and 200 cases was simulated using the CEU allele frequencies to generate control allele counts while the case allele counts were generated using a weighted average of CEU and YRI allele frequencies, with the YRI proportion increasing from 0 to 1. In order to simulate a large study, a sample of 900,000 subjects with equal numbers of cases and controls was simulated with mainly CEU ancestry but with a fraction of 0.01 YRI ancestry in controls and a fraction ranging from 0.01 to 0.011 YRI ancestry in cases.

## Results

Table 1A shows the results of linear regression analysis of the chi-squared for allele frequency differences against LD scores for the LD-pruned SNPs. It can be seen that the LD score is indeed correlated with the difference in allele frequency between CEU and other cohorts. This produces a positive gradient for the regression line and means that the mean chi-squared is higher than the intercept. The effect is most marked in the comparison between CEU and YRI cohorts. The gradient is 0.295 (SE 0.033, p=10^−18^) with an intercept of 42.1 and a mean chi-squared of 46.0. The correlation coefficient between the chi-squared and LD score is 0.0025. Table 1B shows that similar results are obtained for the chromosome 22 SNPs although for some cohorts the correlation is not statistically significant.

**Table 1.**
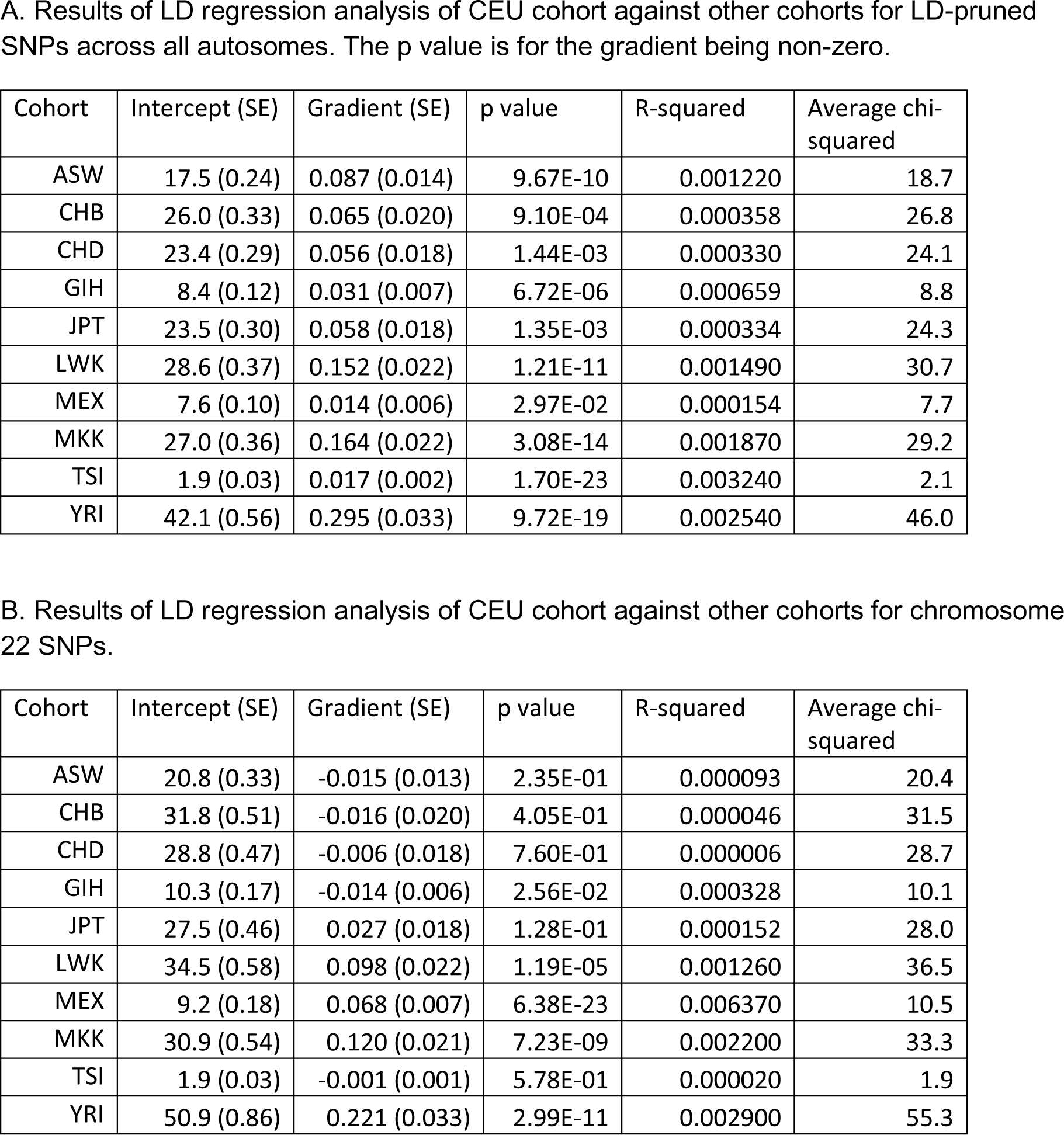

Table 2 shows the results using simulations generated from allele frequencies assuming different proportions of CEU and YRI ancestry. In Table 2A it can be seen that, as would be expected, with large proportions of YRI ancestry in cases the gradient and correlation coefficient increase. However when the proportion of YRI ancestry is less than 0.5 the gradient is very small or even negative, meaning that in this situation the mean chi-squared is equal to or less than the intercept. In Table 2B, intended to reflect a more realistic situation of a large sample size and some YRI ancestry in cases and controls, it can be seen that only a small degree of enrichment of YRI in cases, from 0.01 to 0.0105, is sufficient to increase the intercept to 1.05. With this degree of enrichment the gradient is very small and essentially there is no inflation of the mean chi-squared. When the enrichment increases to 0.011, producing an intercept of 1.25, the gradient becomes very slightly negative, actually producing a mean chi-squared which is slightly smaller than the intercept.

**Table 2.**

| Proportion YRI | Intercept (SE) | Gradient (SE) | p value | R-squared | Average chi-squared |
| --- | --- | --- | --- | --- | --- |
| 0.0 | 1.03 (0.02) | 0.000 (0.001) | 9.73E-01 | 0.000000 | 1.03 |
| 0.1 | 1.93 (0.03) | -0.005 (0.001) | 4.23E-04 | 0.000817 | 1.84 |
| 0.2 | 4.08 (0.07) | -0.007 (0.003) | 1.29E-02 | 0.000406 | 3.94 |
| 0.3 | 7.37 (0.13) | -0.006 (0.005) | 2.27E-01 | 0.000096 | 7.25 |
| 0.4 | 11.40 (0.20) | 0.004 (0.008) | 5.90E-01 | 0.000019 | 11.50 |
| 0.5 | 16.70 (0.28) | 0.010 (0.011) | 3.49E-01 | 0.000058 | 16.90 |
| 0.6 | 22.60 (0.38) | 0.035 (0.015) | 1.54E-02 | 0.000386 | 23.30 |
| 0.7 | 29.60 (0.49) | 0.072 (0.019) | 1.40E-04 | 0.000953 | 31.00 |
| 0.8 | 37.50 (0.62) | 0.117 (0.024) | 1.01E-06 | 0.001570 | 39.80 |
| 0.9 | 47.10 (0.78) | 0.166 (0.030) | 3.47E-08 | 0.002000 | 50.40 |
| 1.0 | 57.40 (0.96) | 0.249 (0.037) | 1.64E-11 | 0.002980 | 62.40 |

| Proportion YRI | Intercept (SE) | Gradient (SE) | p value | R-squared | Average chi-squared |
| --- | --- | --- | --- | --- | --- |
| 0.0100 | 0.99 (0.02) | 0.000 (0.001) | 5.33E-01 | 0.000026 | 1.00 |
| 0.0101 | 1.03 (0.02) | 0.000 (0.001) | 5.48E-01 | 0.000024 | 1.02 |
| 0.0102 | 1.04 (0.02) | -0.001 (0.001) | 2.99E-01 | 0.000071 | 1.03 |
| 0.0103 | 1.02 (0.02) | 0.001 (0.001) | 2.77E-01 | 0.000078 | 1.04 |
| 0.0104 | 1.04 (0.02) | 0.000 (0.001) | 8.26E-01 | 0.000003 | 1.04 |
| 0.0105 | 1.06 (0.02) | 0.000 (0.001) | 5.93E-01 | 0.000019 | 1.05 |
| 0.0106 | 1.11 (0.02) | -0.001 (0.001) | 2.08E-01 | 0.000104 | 1.09 |
| 0.0107 | 1.12 (0.02) | -0.001 (0.001) | 3.13E-01 | 0.000067 | 1.11 |
| 0.0108 | 1.14 (0.02) | -0.001 (0.001) | 3.29E-01 | 0.000063 | 1.12 |
| 0.0109 | 1.17 (0.02) | -0.001 (0.001) | 3.54E-01 | 0.000056 | 1.15 |
| 0.0110 | 1.25 (0.02) | -0.002 (0.001) | 9.18E-03 | 0.000446 | 1.21 |

## Discussion

From these results we draw two main conclusions. The first conclusion is that a fundamental assumption of the LD score regression method, that LD score is not associated with allele frequency differences between populations, is incorrect. The second conclusion is that in practice this does not matter.

When we compare the CEU cohort to others we actually observe quite marked association between the LD score and the chi-squared for allele frequency differences. In the case of the YRI cohort this produces a correlation coefficient ten times higher than that reported between any of the pairs of European cohorts originally studied. This is clearly observed even when LD-pruned SNPs are used, meaning that there can be no artefact for example related to self-correlated SNPs. The positive correlation means that the mean chi-squared is higher than the intercept. However, this is only a small effect. Using the original samples, the mean chi-squared for the YRI cohort is only about 10% higher than the intercept for both the LD-pruned and chromosome 22 SNPs. If this effect scaled linearly with sample size then in practice it would not be expected to produce major problems. However it is not intuitively obvious that this effect would scale linearly and to address this we carried out simulations using a large sample size. What we see is that in fact the effect actually diminishes markedly. When there is only a small degree of enrichment of YRI ancestry, such that the intercept increases from 1 to 1.05 in line with that observed in the association study of risky behaviour, then there is essentially no inflation of the mean chi-squared.

From the investigations we have performed we conclude that the LD score is positively associated with allele frequency differences between populations but that if a low value is observed for the intercept then any increase in the mean chi-squared can be ascribed to a polygenic effect on the phenotype rather than to population stratification.

## Author contributions

MA wrote code and carried out analyses. DC conceived project, carried out further analyses and wrote manuscript.

## Conflict of interest statement

DC was an author on the paper originally describing the LD score regression method. Otherwise, the authors declare no conflict of interest.

## References

Bulik-Sullivan, B. K., Loh, P.-R., Finucane, H. K., Ripke, S., Yang, J., Patterson, N., Daly, M. J., Price, A. L., Neale, B. M. and Neale, B. M. (2015) ‘LD Score regression distinguishes confounding from polygenicity in genome-wide association studies’, Nature Genetics, 47(3), pp. 291–295. doi: 10.1038/ng.3211.

Chang, C. C., Chow, C. C., Tellier, L. C., Vattikuti, S., Purcell, S. M. and Lee, J. J. (2015) ‘Second-generation PLINK: rising to the challenge of larger and richer datasets’, GigaScience. BioMed Central, 4(1), p. 7. doi: 10.1186/s13742-015-0047-8.

Curtis, D. (2018) ‘Polygenic risk score for schizophrenia is more strongly associated with ancestry than with schizophrenia’, Psychiatric Genetics, 28(5), pp. 85–89. doi:10.1097/YPG.0000000000000206.

International HapMap 3 Consortium, D. M., Altshuler, D. M., Gibbs, R. A., Peltonen, L., Altshuler, D. M., Gibbs, R. A., Peltonen, L., Dermitzakis, E., Schaffner, S. F., Yu, F., Peltonen, L., Dermitzakis, E., Bonnen, P. E., Altshuler, D. M., Gibbs, R. A., de Bakker, P. I. W., Deloukas, P., Gabriel, S. B., Gwilliam, R., Hunt, S., Inouye, M., Jia, X., Palotie, A., Parkin, M., Whittaker, P., Yu, F., Chang, K., Hawes, A., Lewis, L. R., Ren, Y., Wheeler, D., Gibbs, R. A., Muzny, D. M., Barnes, C., Darvishi, K., Hurles, M., Korn, J. M., Kristiansson, K., Lee, C., McCarrol, S. A., Nemesh, J., Dermitzakis, E., Keinan, A., Montgomery, S. B., Pollack, S., Price, A. L., Soranzo, N., Bonnen, P. E., Gibbs, R. A., Gonzaga-Jauregui, C., Keinan, A., Price, A. L., Yu, F., Anttila, V., Brodeur, W., Daly, M. J., Leslie, S., McVean, G., Moutsianas, L., Nguyen, H., Schaffner, S. F., Zhang, Q., Ghori, M. J. R., McGinnis, R., McLaren, W., Pollack, S., Price, A. L., Schaffner, S. F., Takeuchi, F., Grossman, S. R., Shlyakhter, I., Hostetter, E. B., Sabeti, P. C., Adebamowo, C. A., Foster, M. W., Gordon, D. R., Licinio, J., Manca, M. C., Marshall, P. A., Matsuda, I., Ngare, D., Wang, V. O., Reddy, D., Rotimi, C. N., Royal, C. D., Sharp, R. R., Zeng, C., Brooks, L. D. and McEwen, J. E. (2010) ‘Integrating common and rare genetic variation in diverse human populations.’, Nature, 467(7311), pp. 52–8. doi: 10.1038/nature09298.

Karlsson Linnér, R., Biroli, P., Kong, E., Meddens, S. F. W., Wedow, R., Fontana, M. A., Lebreton, M., Tino, S. P., Abdellaoui, A., Hammerschlag, A. R., Nivard, M. G., Okbay, A., Rietveld, C. A., Timshel, P. N., Trzaskowski, M., Vlaming, R. de, Zünd, C. L., Bao, Y., Buzdugan, L., Caplin, A. H., Chen, C.-Y., Eibich, P., Fontanillas, P., Gonzalez, J. R., Joshi, P. K., Karhunen, V., Kleinman, A., Levin, R. Z., Lill, C. M., Meddens, G. A., Muntané, G., Sanchez-Roige, S., Rooij, F. J. van, Taskesen, E., Wu, Y., Zhang, F., Auton, A., Boardman, J. D., Clark, D. W., Conlin, A., Dolan, C. C., Fischbacher, U., Groenen, P. J. F., Harris, K. M., Hasler, G., Hofman, A., Ikram, M. A., Jain, S., Karlsson, R., Kessler, R. C., Kooyman, M., MacKillop, J., Männikkö, M., Morcillo-Suarez, C., McQueen, M. B., Schmidt, K. M., Smart, M. C., Sutter, M., Thurik, A. R., Uitterlinden, A. G., White, J., Wit, H. de, Yang, J., Bertram, L., Boomsma, D. I., Esko, T., Fehr, E., Hinds, D. A., Johannesson, M., Kumari, M., Laibson, D., Magnusson, P. K. E., Meyer, M. N., Navarro, A., Palmer, A. A., Pers, T. H., Posthuma, D., Schunk, D., Stein, M. B., Svento, R., Tiemeier, H., Timmers, P. R. H. J., Turley, P., Ursano, R. J., Wagner, G. G., Wilson, J. F., Gratten, J., Lee, J. J., Cesarini, D., Benjamin, D. J., Koellinger, P. D. and Beauchamp, J. P. (2019) ‘Genome-wide association analyses of risk tolerance and risky behaviors in over 1 million individuals identify hundreds of loci and shared genetic influences’, Nature Genetics, 51(2), pp. 245–257. doi: 10.1038/s41588-018-0309-3.

Purcell, S. M., Wray, N. R., Stone, J. L., Visscher, P. M., O’Donovan, M. C., Sullivan, P. F., Sklar, P., Purcell Leader, S. M., Ruderfer, D. M., McQuillin, A., Morris, D. W., O’Dushlaine, C. T., Corvin, A., Holmans, P. a, Macgregor, S., Gurling, H., Blackwood, D. H. R., Craddock, N. J., Gill, M., Hultman, C. M., Kirov, G. K., Lichtenstein, P., Muir, W. J., Owen, M. J., Pato, C. N., Scolnick, E. M., St Clair, D., Sklar Leader, P., Williams, N. M., Georgieva, L., Nikolov, I., Norton, N., Williams, H., Toncheva, D., Milanova, V., Thelander, E. F., Sullivan, P. F., Kenny, E., Quinn, E. M., Choudhury, K., Datta, S., Pimm, J., Thirumalai, S., Puri, V., Krasucki, R., Lawrence, J., Quested, D., Bass, N., Crombie, C., Fraser, G., Leh Kuan, S., Walker, N., McGhee, K. a, Pickard, B., Malloy, P., Maclean, A. W., Van Beck, M., Pato, M. T., Medeiros, H., Middleton, F., Carvalho, C., Morley, C., Fanous, A., Conti, D., Knowles, J. a, Paz Ferreira, C., Macedo, A., Helena Azevedo, M., Kirby, A. N., Ferreira, M. a R., Daly, M. J., Chambert, K., Kuruvilla, F., Gabriel, S. B., Ardlie, K. and Moran, J. L. (2009) ‘Common polygenic variation contributes to risk of schizophrenia and bipolar disorder.’, Nature, 10(AuGuST), pp. 8192–8192. doi: 10.1038/nature08185.

Purcell, S., Neale, B., Todd-Brown, K., Thomas, L., Ferreira, M. A. R., Bender, D., Maller, J., Sklar, P., de Bakker, P. I. W., Daly, M. J. and Sham, P. C. (2007) ‘PLINK: a tool set for whole-genome association and population-based linkage analyses.’, American journal of human genetics. Elsevier, 81(3), pp. 559–75. doi:10.1086/519795.

R Core Team (2014) R: A language and environment for statistical computing. Vienna, Austria., Austria.: R Foundation for Statistical Computing.

Schizophrenia Working Group of the Psychiatric Genomics Consortium (2014) ‘Biological insights from 108 schizophrenia-associated genetic loci’, Nature, 511, pp. 421–427. doi: 10.1038/nature13595.

